# Elucidating the Design Space of Generative Models for Single-Cell Perturbation Prediction

**DOI:** 10.64898/2026.06.15.732063

**Authors:** Sanjukta Bhattacharya, Christian Gensbigler, Shaamil Karim, Jon Lees

## Abstract

Next-token prediction has produced predictable scaling in language, but the recipe presumes a sequence of tokens with a meaningful order. Single-cell RNA-seq counts have no natural gene ordering, so applying the recipe directly to raw expression fails under an ill-suited left-to-right bias. We instead ask whether a learned latent can supply the structure the recipe needs. We introduce `ExpressionVAE` (eVAE), a discrete-latent perturbation model that compresses each cell into a short sequence of discrete codes through a finite-scalar-quantization (FSQ) bottleneck and trains a perturbation-conditioned discrete prior over those codes. On Replogle and Parse 1M, eVAE sets a new state of the art on every distributional metric and leads on most cell-eval perturbation metrics, with Fréchet distance and MMD^2^ roughly 3 to 20× lower than the strongest continuous-latent baseline. Swapping the prior between autoregressive and masked discrete diffusion leaves performance near-identical, isolating the gain to the discrete latent itself rather than the prior family. A decoder-head ablation then exposes a single design axis, the richness of the predictive distribution at inference, that splits the standard metrics into two groups, variance-sensitive and mean-sensitive, which move in opposite directions along the axis. Finally, on a held-out CRISPRi reversion benchmark of 1,732 perturbations under inflammatory cytokine stress, the frozen eVAE encoder outperforms UMAP and differential expression and matches scGPT on perturbation ranking at a fraction of the data.^1^ We release our code.^2^

## 1 Introduction

A cell can be viewed as an information-processing system that maps environmental inputs to molecular responses through an internal regulatory program (10). While the structure of the regulatory network itself is not directly accessible, single-cell RNA sequencing provides a high-dimensional partial observation of its activity (25; 12; 42). Crucially, this observation is discrete: we do not measure continuous expression levels, but count individual molecular events. Across many cells and conditions, these observations implicitly encode the statistical dependencies induced by the underlying regulatory structure. The objective of a virtual cell model is to learn these dependencies well enough to generate realistic cellular states and to predict how modifications to the regulatory program, such as genetic or chemical perturbations, alter the resulting transcriptome (7; 1). If reliable, virtual cells stand to enable a new frontier of in-silico drug discovery.

The central object of observed gene expression is the count matrix: a sparse, zero-inflated, high-dimensional integer-valued tensor recording how many mRNA molecules of each gene were captured from each cell. These properties make direct generative modeling on raw counts remarkably difficult for most models. Autoregressive factorizations impose a sequential ordering on genes, an inductive bias misaligned with their natural exchangeability. Flow matching tends to collapse to the mean under the sparsity of count data. Masked diffusion is the only family that recovers reasonable structure on raw counts (43; 4).

These difficulties on raw counts motivate a latent-space approach. Autoregressive models have demonstrated predictable scaling laws on language data (14), an appealing property as virtual cell models scale up, and they operate naturally over discrete tokens. This motivates a discrete latent representation.

We introduce `ExpressionVAE`, a scalar-quantized variational autoencoder that encodes each cell as a fixed-length sequence of discrete codes, paired with a perturbation-conditioned prior over those codes. Scalar quantization commits the encoder to a finite codebook of cell-state representations that any standard discrete prior can model exactly. To control for the choice of prior, we train two from different generative-model families, autoregressive (AR) and masked discrete diffusion (MDLM); any gain that survives the swap is attributable to the latent representation rather than to a particular prior architecture.

Beyond the latent and prior, two further design questions recur in the single-cell perturbation literature, and we treat each as a controlled experiment. First, single-cell models vary substantially in their decoder output head, and published rankings across cross-entropy, MSE, hurdle, and negative-binomial heads are inconsistent in ways that often track the choice of evaluation metric rather than any property of the heads themselves. We run a within-architecture head ablation on the same trained backbone to expose the underlying axis. Second, we evaluate the frozen encoder on a CRISPRi reversion benchmark to test whether the resulting design choices reflect a generalizable inductive bias rather than over-fitting our chosen metric suite.

Concretely, our contributions are:

- **Discrete-latent perturbation modeling**. We introduce a discrete-latent representation learning approach for single-cell perturbation modeling, and show that pairing `ExpressionVAE` with either an autoregressive or a masked-diffusion prior achieves state-of-the-art on the distributional metrics and leads on most cell-eval state metrics on both Replogle and Parse 1M, with MMD^2^ and Fréchet distance roughly 3–20× lower than the strongest continuous-latent baseline. The two priors achieve essentially identical performance, isolating the gain to the discrete latent representation rather than to the prior architecture (Section 4.1).
- **A measurement-theoretic account of decoder-head choice**. We show that the apparent conflicts between published evaluation metrics resolve into a clean two-group structure once a single design axis, the richness of the inference-time sampling distribution, is made explicit. Every metric is either *variance-sensitive*, rewarding heads that reproduce within-perturbation spread, or *mean-sensitive*, rewarding heads that emit clean per-gene means (Section 4.2).
- **Usefulness of the latent space**. On a held-out CRISPRi reversion benchmark of 1,732 perturbations under inflammatory cytokine stress (41), the hurdle decoder identified by our controlled ablation is the strongest pairing on biological selectivity, and the frozen encoder effectively matches a 10×-larger foundation model (scGPT) on enrichment AUC for perturbation-ranking. This shows that the discrete-latent perturbation prior captures inflammation-axis structure effectively (Section 4.3).

## 2 Related Work

We draw from two literatures: single-cell generative and perturbation-response models, and the latent modeling toolkit from vision and language.

### Generative and perturbation-response models

scVI (18; 19) established the canonical Gaussian-latent, Negative Binomial-likelihood VAE recipe, elaborated by the conditional-VAE thread of scGen, trVAE, and CPA (20; 21; 22) and the latent-diffusion thread of scDiffusion (24), CFGen (30), and scLDM (29) with flow-matching priors (16; 31); the Hurdle head we evaluate traces to MAST (9). Beyond this lineage, perturbation-response models pretrained on tens to hundreds of millions of cells (Geneformer (37), scGPT (6), scFoundation (13), STATE (2), Tahoe-x1 (11)) increasingly condition on ESM-2 embeddings (15). All embed raw counts into continuous vectors. A concurrent line generates directly in the discrete count domain, via Score Entropy Discrete Diffusion (DCM (4; 23)), absorbing-state masked diffusion in the D3PM (3) lineage (Lingshu-Cell (43), X-Cell (39), scDiVa (40)), or Stack’s in-context approach (8). We evaluate on Replogle Perturb-seq (33) and the Parse cytokine atlas (27), with phenotypic reversion (41) as our NF*κ*B ranking experiment.

### Latent generative modeling: tokenizers and priors

On the tokenizer side, the community relies heavily on VQ-VAE as the go-to method for obtaining discrete tokens. VQ-VAE (38; 32) and MaskGIT (5) introduced learnable discrete codes and masked parallel decoding, Latent Diffusion (34) established the continuous-latent plus diffusion-prior recipe, and FSQ (26) eliminated the codebook collapse documented in VQ literature including in single cell domain (17). On the prior side, the absorbing-state masked discrete diffusion family has matured rapidly from D3PM (3) through SEDD (23), MDLM (35), and MD4 (36) to RADD (28), which unifies it with any-order autoregression. We combine learned discrete latents with flexible generative priors, a combination no existing model explores.

## 3 Method

We propose a two-stage pipeline (Fig. 2). Stage 1 trains an `ExpressionVAE` that compresses each cell’s gene-expression vector into a fixed-length sequence of discrete codes via a finite scalar quantization (FSQ) bottleneck (26). Stage 2 freezes the VAE and trains a perturbation-conditioned generative prior over the code sequence, conditioned on an ESM2-3B protein-language-model embedding of the targeted gene and a learned additive cell-type signal. At inference, the prior is queried with the embedding of an unseen perturbation and the resulting latent is decoded into a predicted gene-expression profile. The two design axes varied across the experiments are the VAE *output head* (Section 4.2) and the choice of *prior* (Section 4.1).

**Figure 1:**
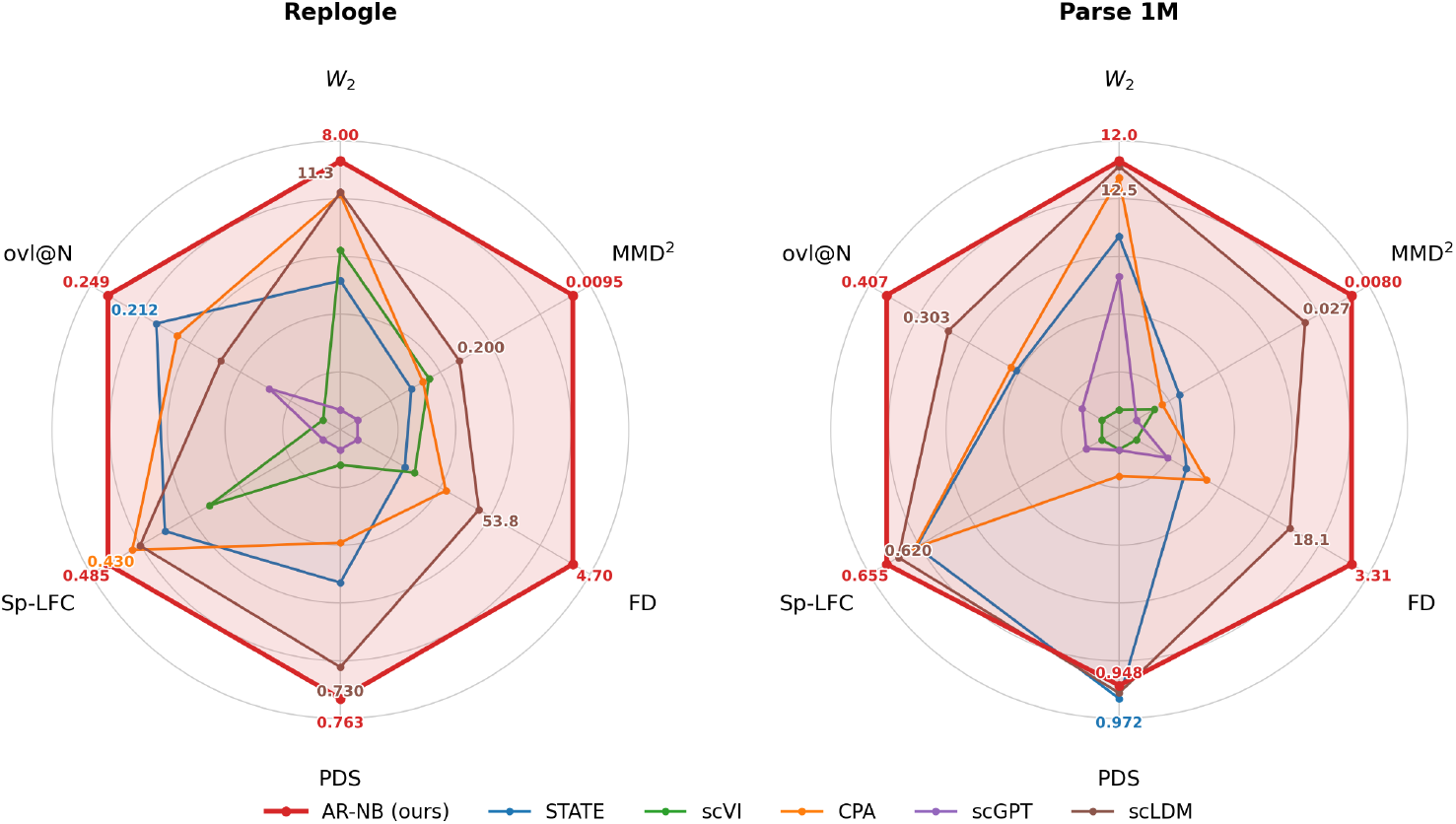
ExpressionVAE on the outer envelope of both metric families. Six-axis comparison of our representative model (Autoregressive prior with Negative Binomial head) against five baselines on Replogle (left) and Parse 1M (right), spanning three distribution metrics (*W*_2_, MMD^2^, FD) and three cell-eval metrics (PDS, Sp-LFC, ovl@N). ExpressionVAE model holds the outermost ring on all six axes on Replogle and five of six on Parse 1M (STATE narrowly leads PDS). It is the only method on the frontier of both metric families: baselines strong on the distribution axes retreat on the ranking axes, and those strong on ranking collapse on the distribution axes. Each axis is rescale and adjusted so that better always points outward.

**Figure 2:**
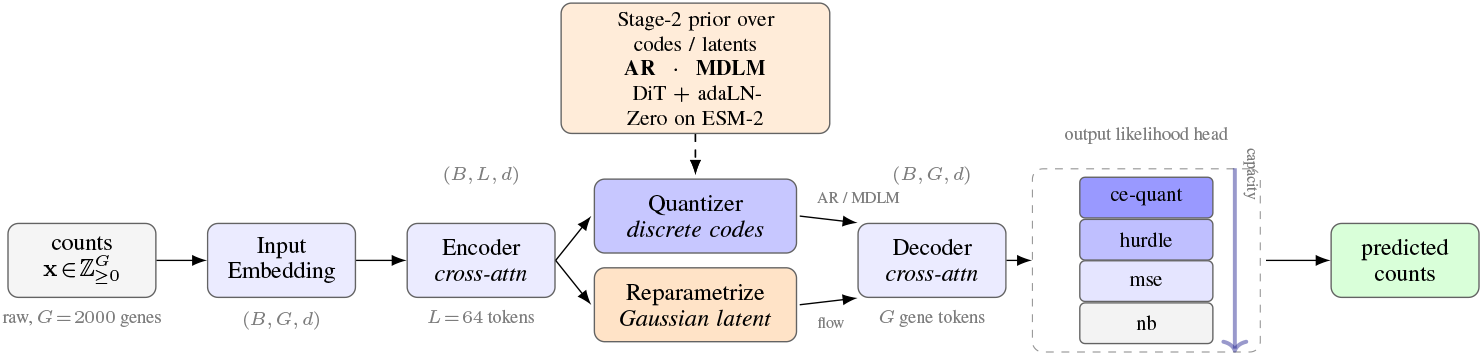
A two-stage pipeline. Stage 1 (the row of boxes): a cross-attention transformer encoder maps raw count vectors 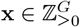 to *L*=64 latent tokens, which are either FSQ quantised into discrete codes (consumed by AR or MDLM priors) or reparametrised as continuous Gaussian latents (consumed by flow). A symmetric cross-attention decoder maps these back to per-gene representations, which one of four output likelihood heads converts to predictions. Stage 2 (sidebar): a DiT prior over the frozen codes / latents, conditioned on the ESM-2 perturbation embedding and optionally cell-type embedding via adaLN-Zero.

### 3.1 Stage 1: ExpressionVAE

#### Encoder and decoder

The encoder maps a count vector to a set of *L* latent tokens via a cross-attention transformer. Each gene *g* is first embedded as **e**_*g*_ = *f*_in_(*x*_*g*_, **v**_*g*_), where *f*_in_ is an MLP applied to the log1p-normalized scalar count *x*_*g*_ and modulated by a learned gene-identity token **v**_*g*_ ∈ ℝ^*d*^. *L* learnable latent query tokens cross-attend over the *G* gene embeddings through a stack of transformer layers, producing a per-cell latent matrix **Z** ∈ ℝ^*L×d*^. A symmetric cross-attention transformer decodes the quantized codes back to gene space: *G* learnable gene query tokens cross-attend over the projected code embeddings, yielding a per-gene representation **H** ∈ ℝ^*G×d*^.

#### FSQ bottleneck

The encoder output **Z** is quantized per-dimension by FSQ (26), which replaces the learned Vector Quantized (VQ) codebook with a parameter-free per-dimension rounding scheme. Given a level vector 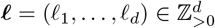, each dimension is bounded via tanh and rounded to the nearest integer on a uniform grid:

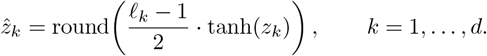

The result is a per-cell sequence of *L* discrete codes **c** = (*c*_1_, …, *c*_*L*_) drawn from a vocabulary of size Π_*k*_ *l*_*k*_, eliminating the codebook collapse typically seen in VQ, also observed in single-cell literature (17). We use *L* = 64 tokens with codebook size *K* = 512. A Gaussian-latent variant used in the Appendix also uses *L* = 64 latent tokens with *β*_KL_ = 2 × 10^*−*6^.

#### Output heads

An output head maps the per-gene representation **H** to a conditional distribution *p*_*θ*_(**x** | **H**). We evaluate four head families, corresponding to standard choices in the single-cell modeling literature, and refer to them by the labels used in Section 4.2.

`ce-quant` places a categorical over [0, *c*_max_] per gene with logits produced by a linear projection of **h**_*g*_, trained against quantile-binned integer targets and sampled at inference (Section 3.1 below).

`hurdle` (9) parameterises a Bernoulli zero-gate *z*_*g*_ ~ Bern(1 − *p*_0,*g*_) together with a regressed positive magnitude *µ*_*g*_; the loss combines BCE on the gate with MSE on non-zero entries, and inference samples the gate while emitting *µ*_*g*_ deterministically.

`mse` regresses log1p-normalised counts directly, with *p*(*x*_*g*_ | **h**_*g*_) = *N* (*µ*_*g*_, 1), and emits *µ*_*g*_ with no stochastic step.

The negative binomial (NB) family parameterises NB(*µ*_*g*_, *θ*_*g*_) per gene with library-size factorisation, followI:ing (18; 29): *µ*_*g*_ = *l*_*n*_ · *ρ*_*g*_, where ***ρ*** = softmax_*g*_(**w**^***⊤***^**H**) is a probability simplex over genes, *l*_*n*_ =**Σ**_*g*_ *x*_*g*_ is the per-cell library size, and *θ*_*g*_ is a learned cell-independent dispersion. This family admits two distinct `predict()` rules at inference time without changing trained weights: `nb-sample` draws from NB(*µ, θ*), and `nb-deter` returns *µ* directly. We treat these as two separate evaluation points throughout, since the choice of inference rule materially shifts which evaluation metrics the head wins (Section 4.2).

#### Quantile binning for the categorical head

The `ce-quant` head requires integer targets, so we quantile-bin the raw counts into a vocabulary of 20 bins chosen from the empirical training count distribution: 8 singleton bins for counts 0–7 capture the dominant low-count regime exactly, 7 bins covering 8–63 handle moderate counts at progressively coarser resolution, and 5 bins (64–95, 96–127, 128–191, 192–255, ≥ 256) cover the high-expression tail. Tokenisation maps each raw count to its bin index; detokenisation maps each predicted bin back to the empirical conditional mean of counts within that bin range, not the arithmetic midpoint, since the within-bin distribution is heavy-tailed and the bias matters most for the unbounded ≥ 256 bucket. This scheme captures 100% of top-DEG per-perturbation means on Parse and 99.69% on Replogle within bounded bins.

### 3.2 Stage 2: Discrete prior over latent codes

Stage 2 freezes the VAE and trains a generative prior over the code sequence **c** = (*c*_1_, …, *c*_*L*_), conditioned on a projected ESM2-3B perturbation embedding **p** and an additive cell-type signal. Both prior families share a DiT backbone with adaLN-Zero conditioning on **p** and the cell-type vector. The autoregressive (AR) prior factorises as 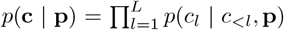 via a causally-masked DiT, requiring *L* sequential decoding steps at inference. MDLM (35) trains a bidirectional DiT denoiser under an absorbing-state masking schedule *α*_*t*_ = 1 − *t*, with the standard masked-token loss

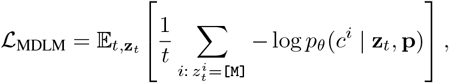

and denoises all *L* tokens in a single forward pass. We use the same backbone, parameter count, and training schedule for both, so any gap in Section 4.1 reflects the prior family rather than capacity.

### 3.3 Inference

Given an unseen perturbation, the corresponding ESM2-3B embedding **p**^*\**^ is passed to the trained prior, which samples a latent sequence ĉ ~ *p*_*θ*_(·| **p**^*\**^) via *L* autoregressive steps for AR or *D* denoising steps for MDLM. The frozen VAE decoder maps **ĉ** back to gene space, and the chosen output head emits a predicted profile 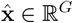 following its `predict()` rule. The same frozen encoder is reused without any fine-tuning for the reversion ranking evaluation in Section 4.3.

## 4 Experiments

### Datasets

We evaluate on two large-scale single-cell perturbation screens. The train and heldout splits are matched to scLDM (29). Parse 1M consists of cytokine perturbations across approximately one million cells, with a CD4 Naive holdout of 27 cytokines. The Replogle essential-genome CRISPRi screen (33) provides a HepG2 holdout of 372 knock-outs.

### Metrics

We evaluate at two complementary levels. *Distribution* metrics (*W*_2_, MMD^2^ with an RBF kernel, and Fréchet distance *FD*), following (29), compare the full predicted cell population against the true cells in PCA space. The *cell-eval* per-perturbation metrics from (2) score the predicted transcriptional change relative to control: *PR-AUC* and *Spearman sig* are built on FDR-based DE-test calls, PDS measures perturbation specificity from the pseudobulk effect vector, *Spearman LFC* and *overlap@N* rank genes by predicted log_2_ fold change, and *Pearson* Δ correlates the full per-gene Δ vector. All numbers report mean ± SE across four seeds, with full definitions in Appendix C.

### 4.1 Discrete latents outperform continuous baselines

Existing single-cell perturbation models build on continuous latent representations (scVI, CPA, scLDM) or operate directly in expression space (scGPT, STATE). Discrete-latent representation learning, despite being the foundation of the strongest generative models in language and vision, has not previously been applied to perturbation modeling. We close this gap. The `ExpressionVAE` encodes each cell as a fixed-length sequence of discrete codes drawn from a scalar-quantized bottleneck, and a perturbation-conditioned prior, either autoregressive (AR) or masked discrete diffusion (MDLM), models the joint distribution over those codes. We compare against six baselines spanning continuous latent VAEs, conditional perturbation models, expression-space foundation models, and a non-parametric mean baseline: scLDM (29), scVI (18), CPA (22), scGPT (6), STATE (2), and PerturbMean. The results of Replogle and Parse 1M are reported in Table 1.

**Table 1:** Discrete-latent perturbation modeling against six baselines on Replogle and Parse 1M. For each prior (AR, MDLM) we report two rows, corresponding to the NB-sample and MSE output heads (Section 4.2). Bold = best in column for that dataset, * = scores are re-used from original reports.

| Dataset | Prior | $W_2 \downarrow$ | $\text{MMD}^2 \downarrow$ | $\text{FD} \downarrow$ | $\text{PDS} \uparrow$ | $\text{PR-AUC}^* \uparrow$ | $\text{Sp-sig}^* \uparrow$ | $\text{Sp-LFC} \uparrow$ | $\text{ovl@N} \uparrow$ | $\text{P-}\Delta^* \uparrow$ |
| --- | --- | --- | --- | --- | --- | --- | --- | --- | --- | --- |
| Replogle | AR | **8.001** | **0.009** | **4.70** | **0.763** | 0.295 | 0.789 | 0.480 | 0.170 | 0.307 |
|  |  | 9.032 | 0.099 | 35.06 | 0.716 | 0.230 | 0.793 | 0.508 | **0.255** | 0.381 |
|  | MDLM | 8.121 | 0.013 | 5.29 | 0.756 | **0.297** | **0.799** | 0.495 | 0.192 | 0.318 |
|  |  | 9.091 | 0.105 | 35.70 | 0.698 | 0.219 | 0.793 | **0.540** | 0.248 | 0.362 |
|  | STATE | 20.580 | 0.730 | 366.64 | 0.641 | 0.210 | 0.721 | 0.357 | 0.212 | **0.420** |
|  | ScVI | 17.359 | 0.453 | 284.47 | 0.517 | 0.190 | 0.598 | 0.257 | 0.085 | 0.350 |
|  | CPA | 11.510 | 0.532 | 126.81 | 0.599 | 0.190 | 0.370 | 0.430 | 0.196 | 0.380 |
|  | ScGPT | 34.166 | 3.087 | 1247.68 | 0.501 | 0.100 | 0.550 | 0.002 | 0.126 | 0.385 |
| | ScLDM ($\omega = 1$) | 11.292 | 0.200 | 53.75 | 0.730 | — | — | 0.413 | 0.163 | — |
|  | PerturbMean | — | — | — | 0.616 | 0.190 | 0.190 | 0.246 | 0.162 | — |
| Parse 1M | AR | **11.952** | **0.008** | **3.31** | 0.948 | **0.498** | 0.949 | **0.655** | **0.407** | 0.762 |
|  |  | 22.080 | 0.646 | 375.41 | 0.632 | 0.157 | 0.926 | 0.618 | 0.140 | 0.747 |
|  | MDLM | 11.969 | 0.008 | 3.50 | 0.956 | 0.485 | **0.960** | 0.639 | 0.401 | **0.763** |
|  |  | 22.152 | 0.654 | 378.10 | 0.624 | 0.141 | 0.901 | 0.646 | 0.144 | 0.733 |
|  | STATE | 19.111 | 0.714 | 312.34 | 0.972 | — | — | 0.565 | 0.188 | — |
|  | ScVI | 35.508 | 1.372 | 1233.11 | 0.519 | — | — | 0.010 | 0.044 | — |
|  | CPA | 13.534 | 1.117 | 181.32 | 0.567 | — | — | 0.570 | 0.197 | — |
|  | ScGPT | 22.870 | 2.203 | 523.93 | 0.520 | — | — | 0.057 | 0.077 | — |
| | ScLDM ($\omega = 1$) | 12.457 | 0.027 | 18.14 | 0.962 | — | — | 0.620 | 0.303 | — |
|  | PerturbMean | — | — | — | **0.973** | — | — | 0.467 | 0.118 | — |

For each prior we report two rows, corresponding to the NB-sample and MSE output heads. These sit at the sampling and deterministic ends of the head spectrum analyzed in Section 4.2 and trade off across metric families; the practical takeaway here is that our model achieves the strongest result in each family. Three observations stand out.

#### 3–20× gains on distributional metrics

The headline result is on *W*_2_, MMD^2^, and FD, where the discrete-latent model improves over the strongest continuous-latent baseline (scLDM) by ~10–20× on Replogle and ~3–6× on Parse 1M, on MMD^2^ and FD. On Replogle, MMD^2^ drops from 0.200 to 0.009 and FD from 53.75 to 4.70; on Parse 1M, MMD^2^ drops from 0.027 to 0.008 and FD from 18.14 to 3.31. Expression-space methods (scGPT, STATE) and continuous latent baselines (scVI, CPA) lag the discrete-latent model by one to three orders of magnitude on these metrics.

#### Two priors, same numbers

AR and MDLM, drawn from different generative-model families and commonly viewed as alternatives, achieve effectively identical performance across all metrics on both datasets. This is direct evidence that the gain is from the discrete-latent representation rather than from a particular prior architecture, since the two priors share only the encoder and the codebook.

#### Top of both metric families

The model achieves the strongest result on every distributional metric and most DE-discovery metrics *and* on every mean-ranking metric, on both datasets. No baseline reaches the top of both families. Existing methods are competitive on distributional metrics or on ranking metrics, but never both: scLDM is the strongest baseline on distributional metrics yet trails on *Sp-LFC* and *overlap@N*, while PerturbMean and STATE are competitive on PDS and mean-ranking metrics but collapse on *W*_2_, MMD^2^, and FD. The two-rows-per-prior structure that produces this pattern is not incidental; Section 4.2 shows it reflects a single underlying axis governed by inference-time stochasticity.

### 4.2 Sampling-richness axis explains decoder-head behavior

A practical question for any single-cell perturbation model is which decoder head to use: cross-entropy, hurdle, MSE, negative binomial. The literature offers no consistent answer. Different papers report different metrics, and different metrics rank these heads differently. The two rows per prior in Table 1 make the conflict visible within a single backbone: the same trained model produces noticeably different metric profiles depending only on the head. We show here that this is not noise. The standard evaluation metrics partition into two groups, and within each group the head ordering is monotonic in a single underlying axis, the richness of the inference-time sampling distribution, with the two groups responding in opposite directions along it.

#### Five heads on a sampling-richness gradient

We evaluate five heads on the same trained `ExpressionVAE` backbone. `ce-quant` parameterizes a categorical over [0, *c*_max_] per gene and samples from the softmax. `nb-sample` parameterizes a negative binomial NB(*µ, θ*) per gene and emits a draw from it. `hurdle` parameterizes a Bernoulli zero-gate combined with a deterministic positive magnitude *µ*, sampling only the gate. `mse` emits a deterministic real-valued *µ* everywhere. `nb-deter` is just an NB head without sampling, it deterministically returns *µ*. Read in the order `ce-quant` → `nb-sample` → `hurdle` → `mse` → `nb-deter`, the heads form a gradient in the richness of the inference-time predictive distribution: full count sampling, parametric count sampling, sampled zero-gate over deterministic magnitude, fully deterministic, and fully deterministic plus structurally anti-sparse (due to softmax over genes in NB).

#### Two groups, opposing trends

Figure 3 reports all five heads under both priors as per-metric heatmaps, with columns ordered so that the variance-sensitive group precedes the mean-sensitive group. The structure is immediate from the color gradient. The variance-sensitive columns (all three distribution metrics plus DE-discovery: *disc_l1, PR-AUC, Spearman sig*) darken toward the sampling end of the head ordering, the top of each prior block; the mean-sensitive columns (*Spearman LFC, overlap@N, Pearson* Δ) darken toward the deterministic end, the bottom. The variance-sensitive collapse is most dramatic on Parse 1M, where moving the head from sampling to deterministic shifts FD by more than two orders of magnitude (e.g., 1.84 →454.1 along the AR block); on Replogle the same trend holds in direction but is compressed in magnitude, consistent with that dataset’s lower count dispersion.

**Figure 3:**
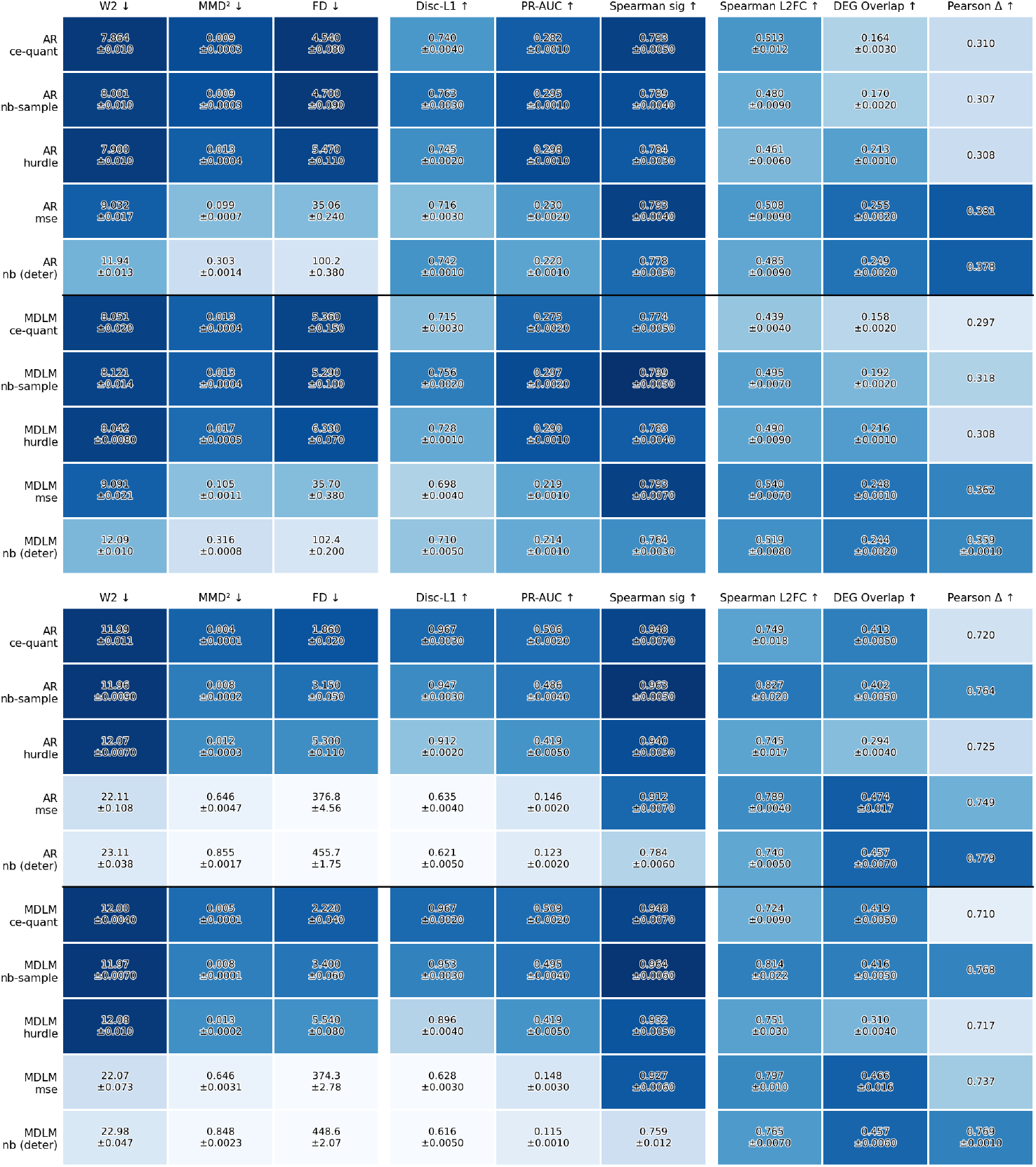
Output-head ablation on Replogle (top) and Parse 1M (bottom). Rows are five output heads under AR and MDLM priors on the same trained backbone; columns are grouped into the two functional families, variance-sensitive (distribution and DE-discovery) and mean-sensitive (mean-ranking). Cells shaded per column, darker indicates better; mean over four seeds, SE in ± notation. Parse 1M DE-based metrics in this figure use a DE-significance threshold of *ϵ* = 0.5.

**Figure 4:**
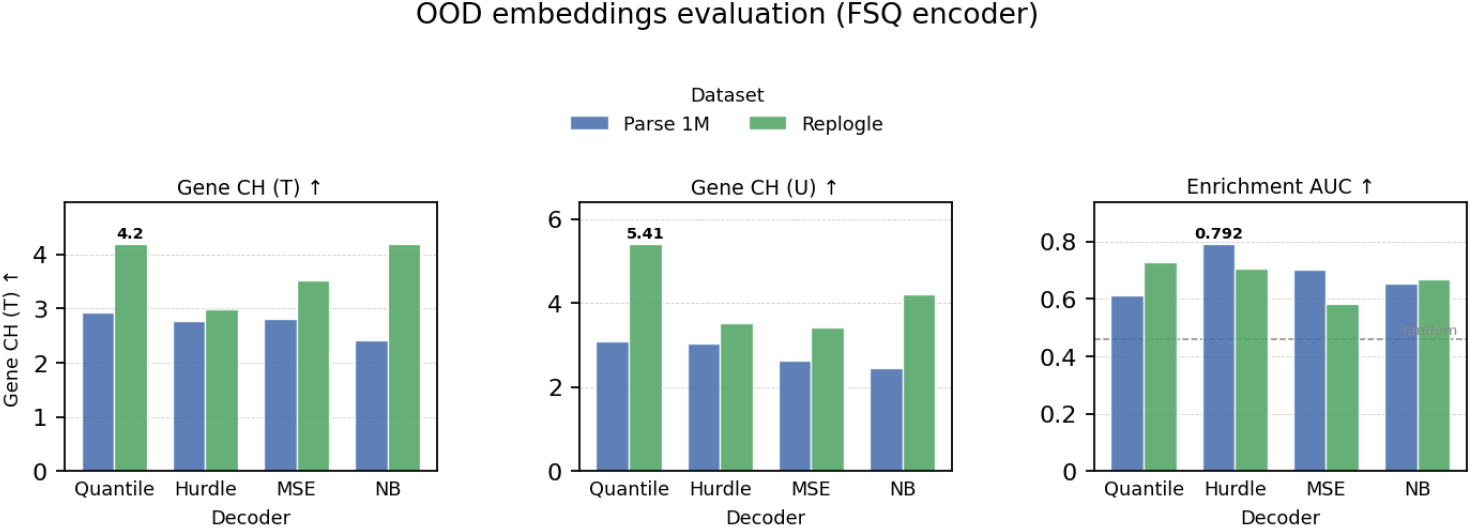
Latent space evaluation on the TeloHAEC phenotypic-reversion benchmark. Frozen `ExpressionVAE` encoders trained on Replogle and Parse 1M. Dashed line on the right shows the random baseline (0.46); annotated values mark the best configuration per panel.

**Figure 5:**
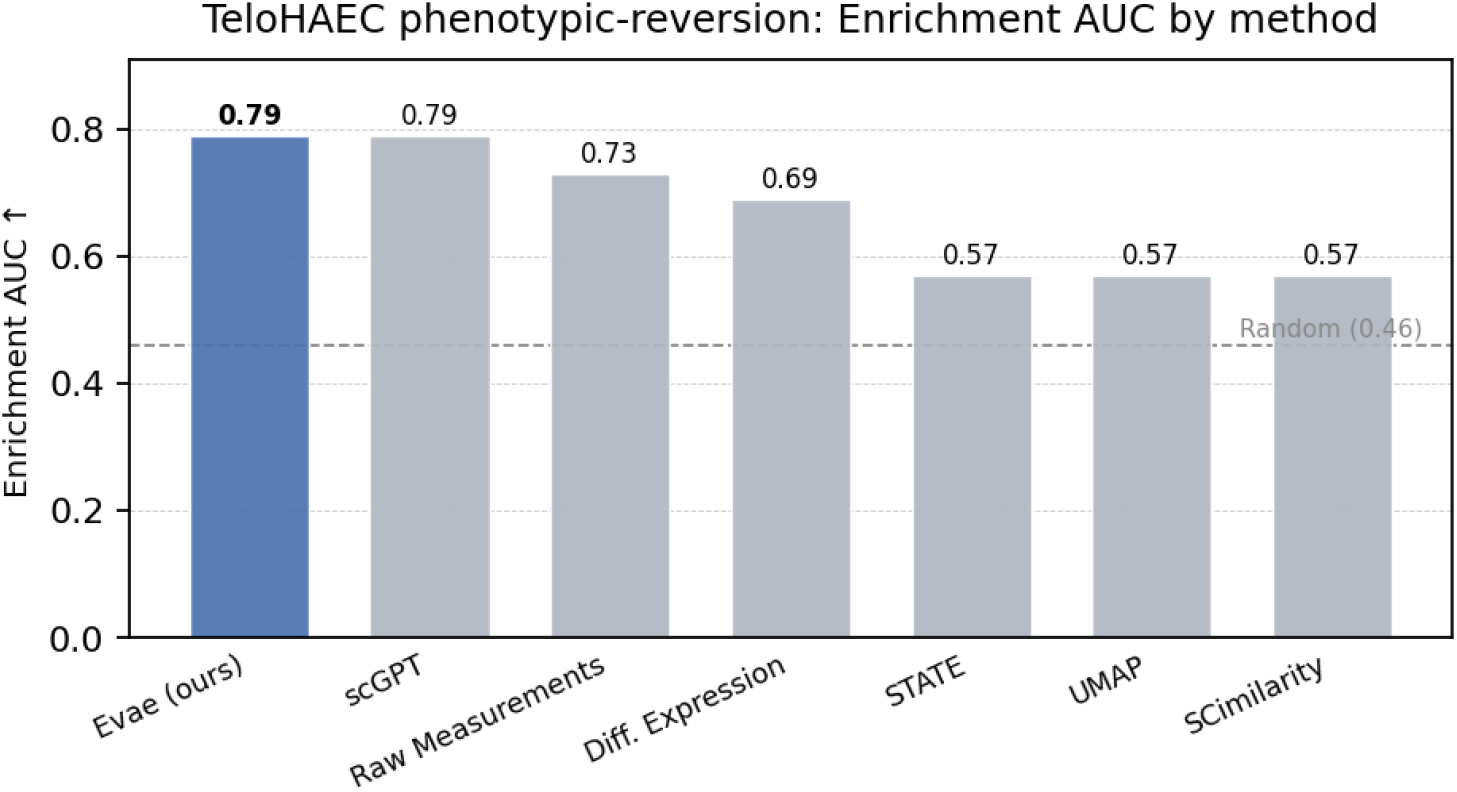
Enrichment AUC for the TeloHAEC phenotypic-reversion benchmark. Dashed line on the right shows the random baseline (0.46); annotated values mark the best configuration per panel.

The mechanism follows from how each metric is computed (Appendix C). The prior draws a fresh latent per cell (Section 3.3), so the predicted population is already spread across cell states for every head; what the head controls is the conditional count noise at a fixed latent. A sampling head adds the per-gene over-dispersion and zero-inflation of real counts on top of each latent’s mean, whereas a deterministic head returns the mean, leaving an *under-dispersed* population. The variance-sensitive metrics penalise this. Fréchet distance, MMD^2^, and *W*_2_ rise as the predicted spread falls short of the true cells; *PR-AUC* and *Spearman sig* compute per-gene FDRs from a Mann–Whitney *U* test that, with too little predicted-side spread it saturates almost any significant mean shift and stops ranking genes by perturbation strength; and PDS sums an *L*_1_ over the full per-gene effect vector, where the under-dispersed, zero-poor pseudobulk is biased across the many low-count genes. The mean-sensitive metrics instead concentrate on the strongly-perturbed genes, where the deterministic mean is cleanest and count noise only adds finite-sample variance: *Spearman LFC* and *overlap@N* are rank-based on the real-DE set, and *Pearson* Δ correlates the full Δ vector but, as a correlation, is dominated by the large-magnitude genes. PDS and *Pearson* Δ make the split concrete: both read the full per-gene vector, but *L*_1_ weights the sparse bulk (variance-sensitive) while correlation weights the peaks (mean-sensitive).

#### The NB pair isolates the sampling axis

A natural confound is that `ce-quant` differs from `mse` on multiple axes simultaneously: discreteness, parametric form, output dimension, and training loss. To isolate sampling versus deterministic prediction from these confounds, we exploit that `nb-sample` and `nb-deter` share trained weights and differ only in `predict()`. Switching from *µ* to a draw from NB(*µ, θ*) moves NB from the bottom tier to the top tier of the variance-sensitive group (e.g., on Parse, FD : 454.06 → 3.31 and PR-AUC : 0.129 → 0.498 under AR), and reverses the direction on the mean-sensitive group (e.g., on Replogle, overlap@N : 0.281 → 0.170 under AR). The shift is largest on the distribution metrics and milder elsewhere, but the direction is consistent within each group. Because no other component of the model changes, the entire effect is localized to the sampling step at inference. This rules out the confounds and pins the cross-group structure on the sampling axis.

#### What metric measures what, and a recommendation

The variance-sensitive group rewards recovery of within-perturbation cell variance, expressed once directly at the cell-distribution level (the distribution metrics) and once filtered through DE testing (the DE-discovery metrics). The mean-sensitive group rewards clean per-gene mean estimates and is, by construction, penalised by sampling noise. The practical implication is that head choice is governed by what the practitioner intends to measure. We recommend reporting at least one metric from each group, and selecting the head to match the downstream task: a sampling head (`ce-quant` or sampled NB) for use cases that consume predicted cell distributions (synthetic-cohort DE, virtual screens, power calculations), and a deterministic-mean head (`mse` or NB-mean) for gene-list ranking against a reference. Reporting any single metric in isolation, or comparing models that report metrics from different groups, is misleading.

### 4.3 Usefulness of latent space for phenotypic reversion

Sections 4.1 and 4.2 establish that a discrete-latent perturbation model with the right decoder head dominates within-distribution metrics, and that head choice is governed by an inference-time sampling axis with a clean metric-to-head map. To test whether these design choices reflect a generalizable inductive bias rather than over-fitting the chosen metric set, we evaluate the *frozen* encoder on a biological discovery task. No information about the biological task’s dataset enters training; the encoder is used only to embed cells, and metrics are computed directly on the resulting latents.

#### Benchmark

The TeloHAEC phenotypic-reversion benchmark (41) is a CRISPRi screen of 1,732 perturbations across 864,115 cells under IL-1*β* and TNF-*α* inflammation. The task is to rank perturbations capable of reverting the inflammatory state, with the canonical NF-*κ*B activators (TNFRSF1A, TRADD, JUNB, JUND, NFKB1, NFKB2) as positive controls. We report three metrics on the frozen latents: per-perturbation Calinski-Harabasz scores among treated cells (Gene CH (T)) and untreated cells (Gene CH (U)), measuring latent separability; and Enrichment AUC, the rank-AUC of the NF-*κ*B positive controls within the cosine-distance reversion ordering of the 1,732 perturbation centroids (random baseline 0.46).

#### The decoder transfers; the encoder pairing is dataset-dependent

The Hurdle decoder identified by the controlled ablations is the strongest pairing on Enrichment AUC on Parse 1M dataset (0.792); the decoder choice transfers cleanly without fine-tuning. Both axes contribute measurably to performance, confirming that the decoder is not just a likelihood-modeling decision but shapes the representation itself.

#### Quantiled inputs dominate separability

On the per-perturbation Calinski-Harabasz scores, FSQ + Quantile is the strongest configuration on both Gene CH (T) and Gene CH (U) (4.20, 5.41 on Replogle). Hard binning in inputs commits the encoder to a finite codebook of phenotypic states, producing sharper per-perturbation cluster structure in the latent space. This is consistent with the distributional-metric gap reported in Section 4.1. Overall, discrete latents allocate representational capacity in a way that preserves cell-state distinctions both within and outside the training distribution.

#### Comparison with scGPT

On the same benchmark, scGPT (6) achieves Enrichment AUC = 0.79. Our strongest Parse-trained configuration reaches 0.79, within typical evaluation noise of scGPT despite scGPT consuming roughly 10× more pre-training data. The point is not a SOTA claim on the TeloHAEC benchmark, but a representation-efficiency claim: a discrete-latent perturbation prior trained on a single 1M-cell dataset effectively matches a foundation model trained on tens of millions of cells, on a biologically meaningful ranking task. The explicit zero-gate decoder identified by our ablation transfers cleanly off the training distribution. We acknowledge that this benchmark has several distribution shifts such as cell type, perturbation set, biological context, etc, making it hard to identify which axis of generalization the model actually uses with the limited training dataset.

## 5 Conclusion

We introduced `ExpressionVAE`, a discrete-latent perturbation model for single-cell data, and showed that pairing a scalar-quantized bottleneck with either an autoregressive or a masked-diffusion prior achieves state-of-the-art on distributional metrics and leads on most cell-eval metrics on Replogle and Parse 1M. Both priors achieve effectively identical performance, isolating the gain to the discrete latent rather than to a particular prior. A controlled output-head ablation further showed that decoder-head choice in this setting is governed by a single inference-time design axis, sampling versus deterministic prediction, with standard evaluation metrics partitioning into two groups along it, variance-sensitive and mean-sensitive; this resolves the apparent conflicts between published metric rankings into a clean two-way mapping between heads and downstream tasks. The hurdle decoder identified by the ablation transfers as the strongest selectivity pairing on a held-out CRISPRi reversion benchmark, where the frozen encoder effectively matches a 10× larger foundation model at a fraction of the training data. Taken together, these results suggest that a compact discrete latent captures perturbation-axis structure effectively and is a promising step toward virtual cells for in-silico discovery.

## A Hyperparameters and training configuration

Tables 2 and 3 list the hyperparameters used for the main-paper VAE backbone and prior models. The same configuration is shared across Parse 1M and Replogle, with two dataset-specific differences: (i) the perturbation conditioning source — ESM2-3B protein-language-model embeddings keyed by the knocked-out gene on Replogle, and a learnable per-cytokine table on Parse 1M (cytokine ligands have no direct protein-sequence analog) — and (ii) the train/holdout split definitions, matched to scLDM in both cases. Both datasets use additive cell-type conditioning on the adaLN-Zero path.

**Table 2:** Stage 1: ExpressionVAE hyperparameters. Shared across both datasets and all four output heads. The Gaussian-latent variant referenced in Section 4.3 swaps the FSQ bottleneck for a Gaussian-posterior bottleneck with the parameters in the lower block; all other settings are unchanged.

|  |  |
| --- | --- |
| Latent sequence length $L$ | 64 |
| FSQ codebook size $K$ | 512 |
| Encoder / decoder | Cross-attention transformer |
| Gene input embedding | MLP over $\log_1 p$ count + learned gene-identity tokens |
| Optimiser | AdamW |
| Learning rate | $6 \times 10^{-4}$ |
| Schedule | Cosine |
| Warm-up steps | 2,500 |
| Training iterations | 75,000 |
| Batch size | 128 |
| Gaussian-latent variant: per-token latent dim | 8 |
| Gaussian-latent variant: $\beta_{\text{KL}}$ | $2 \times 10^{-6}$ |

**Table 3:** Stage 2: prior hyperparameters. Shared across the autoregressive (AR) and masked-discrete-diffusion (MDLM) priors. The DiT backbone is identical (same depth, width, head count, parameter budget) so any gap in Section 4.1 reflects the prior family rather than capacity.

|  |  |
| --- | --- |
| Backbone | DiT (Diffusion Transformer) |
| Conditioning | adaLN-Zero on perturbation + cell-type embedding |
| Model dimension $d_{\text{model}}$ | 384 |
| Attention heads | 8 |
| Layers | 6 |
| Feed-forward dimension | 1,536 |
| Optimiser | AdamW |
| Learning rate | $2 \times 10^{-4}$ |
| Schedule | Cosine |
| Warm-up steps | 3,000 |
| Gradient clipping (max-norm) | 1.0 |
| Training iterations | 75,000 |
| Batch size | 128 |
| MDLM: masking schedule | absorbing, $\alpha_t = 1 - t$ |
| MDLM: $t_\epsilon$ | $5 \times 10^{-3}$ |
| MDLM: inference denoising steps | 32 |

## B Codebook size and latent-token-count ablation

We swept the FSQ latent sequence length *L* ∈ {64, 128} against the codebook size *K* ∈ {512, 1024} on the Replogle (HepG2) holdout, paired with the AR-FSQ-MSE and MDLM-FSQ-MSE configurations from the main paper. We additionally report AR-FSQ-NB, MDLM-FSQ-NB, and a Flow-Gaussian-MSE configuration as neighbouring anchor points in the design space. Results are in Table 4.

**Table 4:** Codebook size and latent-token-count ablation on the Replogle (HepG2) holdout. Bold = best per column.

| Run | PDS ↑ | ovl@N ↑ | PR-AUC ↑ | Sp-sig ↑ | $W_2$ ↓ | MMD<sup>2</sup> ↓ | FD ↓ |
| --- | --- | --- | --- | --- | --- | --- | --- |
| AR-FSQ-MSE $L=128, K=1024$ | 0.7373 | 0.2800 | 0.2225 | 0.7519 | **10.02** | **0.0706** | 56.26 |
| AR-FSQ-MSE $L=128, K=512$ | 0.7398 | **0.2914** | **0.2301** | 0.7499 | 10.17 | 0.0734 | 57.28 |
| AR-FSQ-MSE $L=64, K=1024$ | 0.7474 | 0.2773 | 0.2196 | 0.7563 | 10.06 | 0.0720 | 56.70 |
| AR-FSQ-MSE $L=64, K=512$ | 0.7510 | 0.2834 | 0.2177 | **0.7856** | 10.06 | 0.0716 | 57.19 |
| AR-FSQ-NB | **0.7594** | 0.2790 | 0.2224 | 0.7678 | 12.35 | 0.1514 | 116.30 |
| MDLM-FSQ-MSE $L=128, K=1024$ | 0.7030 | 0.2750 | 0.2071 | 0.7265 | 10.18 | 0.0791 | 59.31 |
| MDLM-FSQ-MSE $L=128, K=512$ | 0.7147 | 0.2790 | 0.2135 | 0.7440 | 10.25 | 0.0776 | 57.75 |
| MDLM-FSQ-MSE $L=64, K=1024$ | 0.7126 | 0.2692 | 0.2064 | 0.7495 | 10.19 | 0.0771 | 57.80 |
| MDLM-FSQ-MSE $L=64, K=512$ | 0.7169 | 0.2816 | 0.2161 | 0.7344 | 10.27 | 0.0777 | 60.17 |
| MDLM-FSQ-NB | 0.7259 | 0.2728 | 0.2112 | 0.7406 | 12.49 | 0.1580 | 118.57 |
| Flow-Gaussian-MSE | 0.7514 | 0.2820 | 0.2204 | 0.7290 | 10.17 | 0.0707 | **55.10** |

The numbers in Table 4 were obtained on a different gene-set than the main-paper results and are therefore not directly comparable to Table 1; they are, however, internally consistent across rows. Based on this ablation we carried forward *L* = 64 and *K* = 512 to the main experiments.

## C Analysis of Evaluation Metrics

We evaluate perturbation prediction across two complementary families of metrics: *differential expression (DE) metrics*, which assess how well a model reproduces the statistical signature of transcriptional change, and *distribution metrics*, which assess the fidelity of the full predicted cell population. We find that certain design choices lead to some statistical metrics generating NaN values for some perturbations. In Table 1 we ignore these NaN values or take them as 0.

To ensure fair comparison, we use the same training/test split as scLDM (29). Metrics like PR-AUC, Sp-sig, and P-Δ are not reported by (29) so we use the values reported as is from (2). We also note that the latter train/test split is not matched in this case.

In the following, we describe each metric, its mathematical definition.

### C.1 Differential Expression Metrics

All DE based metrics share a common upstream computation: for each perturbation *p* we derive a *pseudobulk* summary, compared against the control population of that perturbation.

#### Pseudobulk aggregation

Given a matrix of log_1p_-normalised expression values 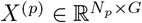 (cells ×genes) for perturbation *p*, and the analogous control matrix 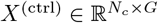, we compute the geometric pseudobulk in count space as

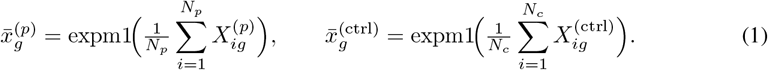

Taking the mean in log space and back-transforming via expm1(*z*) = *e*^*z*^−1 is equivalent to computing the geometric mean of the original (pre-log) counts.

#### Log_**2**_ **fold-change**

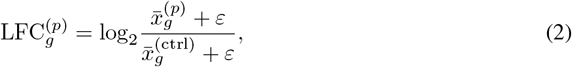

where *ε* = 0 by default; genes with a zero control mean therefore yield ±∞ and are treated as such throughout the pipeline.

#### Per-perturbation FDR

Differential expression is tested per gene with a two-sided Mann–Whitney *U* test applied to the raw (log_1p_) cell vectors of perturbation *p* and controls. *p*-values are corrected to false discovery rates (FDR) using the Benjamini–Hochberg (BH) procedure, applied *independently within each perturbation* (i.e. across the *G* gene *p*-values for that perturbation).

#### C.1.1 Precision–Recall AUC (PR-AUC)

PR-AUC quantifies how well the *predicted* FDR scores rank the truly DE genes. For each perturbation

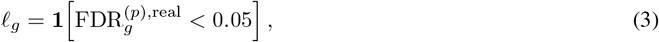

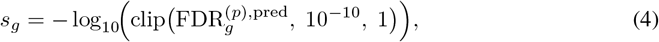

and the per-perturbation area under the precision–recall curve (AP) is computed with `sklearn.metrics.average_precision_score`(***l*, s**). Missing predicted FDR values are filled with 1.0 (least confident), and we clip to [10^*−*10^, 1] before taking the log to avoid numerical issues. We report the mean AP across perturbations.

#### C.1.2 Spearman of Significant-Gene Counts (Sp-sig)

Sp-sig measures across-perturbation agreement in how many genes are called DE, rather than their identities or magnitudes. Let 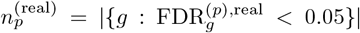 and 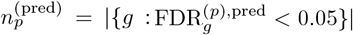. Then

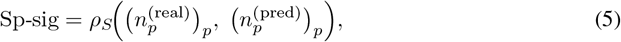

where *ρ*_*S*_ denotes the Spearman rank correlation computed over all perturbations. A model that produces zero significant genes everywhere yields a constant predicted vector; we follow the convention of returning NaN in this degenerate case and use `nanmean` when averaging over seeds.

#### C.1.3 Spearman of Log_2_FC over Significant Genes (Sp-LFC)

Sp-LFC measures per-perturbation agreement in the *direction and magnitude* of DE, restricted to genes that are truly significant. For each perturbation *p*, let 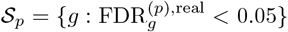 be the real-significant gene set.

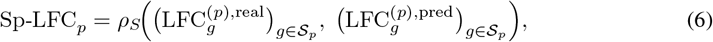

where predicted LFC values for genes absent from the predicted DE table are filled with 0.0. We report the mean over perturbations. Note that both the gene set *S*_*p*_ and the LFC values are defined solely from *real* data; the predicted LFC is only evaluated at those positions.

#### C.1.4 Perturbation Discrimination Score (PDS)

The PDS measures the *specificity* of predicted perturbation effects: does the predicted effect for perturbation *p* match the real effect of *p* more closely than the real effects of all other perturbations? Let 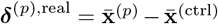 and 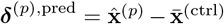 denote the real and predicted perturbation effects (pseudobulk deltas from control). For each perturbation *p*, we compute the *L*_1_ distance between the predicted effect and every real effect:

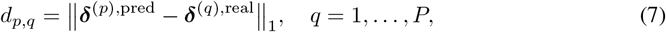

then rank the *P* distances in ascending order and find the rank *r*_*p*_ of the true perturbation *q* = *p*.

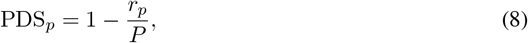

so PDS_*p*_ = 1 indicates the correct perturbation is the nearest neighbour, and PDS_*p*_ = 0.5 is chance performance.

#### CRISPR vs. non-CRISPR data

For CRISPR knockdown experiments, the target gene itself exhibits a strong direct knockdown signal (~ 95% reduction) that would trivially dominate the *L*_1_ distance and inflate PDS for any model that simply predicts a reduction at the target locus. We therefore exclude the target gene from the distance calculation when evaluating on CRISPR perturbation data, following Adduri et al. (2):

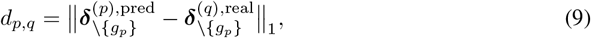

where *g*_*p*_ denotes the target gene of perturbation *p* and the subscript \{*g*_*p*_} denotes the remaining gene dimensions. For non-CRISPR perturbations (cytokines, small molecules) there is no analogous single-gene confound, so no gene is excluded.

### C.2 Distribution-Level Metrics

Beyond DE summary statistics, we evaluate fidelity of the full predicted cell distribution against ground-truth cells for each perturbation. All distribution metrics operate in the top-30 PCA components of the gene expression space; the PCA basis is fitted on the ground-truth cells and used to project both real and predicted populations. We match this to the design details mentioned in (29).

#### C.2.1 Maximum Mean Discrepancy (MMD^2^)

The squared MMD between predicted cells 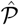 and real cells *Q* under an RBF kernel *k*(**x, y**) = exp(−∥**x** − **y**∥^2^*/σ*^2^) is

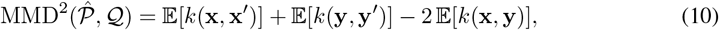

where 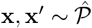 and **y, y**^*′*^ ~ *Q*. We use the median heuristic to set *σ* on the real cells.

#### C.2.2 2-Wasserstein Distance (W_2_)

The 2-Wasserstein distance is defined as the minimum-cost coupling between the two distributions under the squared-*L*_2_ ground metric:

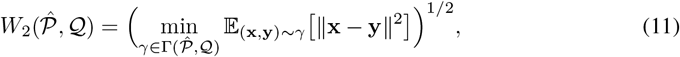

where 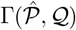 is the set of all couplings. We solve the discrete OT problem exactly via the Earth Mover’s Distance (EMD) solver on subsampled populations (up to 5 000 cells), and report *W*_2_ in PCA-30 space. Sinkhorn is an approximate version of the EMD solver.

#### C.2.3 Fréchet Distance (FD)

The Fréchet distance treats each distribution as a multivariate Gaussian and measures the distance between the fitted Gaussians:

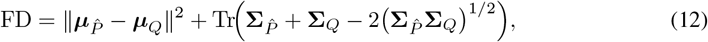

where (·)^1*/*2^ denotes the matrix square root. We add *ε***I** (*ε* = 10^*−*6^) to each covariance matrix before computing the matrix square root to handle rank-deficient PCA components.

### C.3 Implementation Notes and Cross-Library Validation

Table 5 summarises the metrics used in our main evaluation, grouped by the aspect of perturbation response they assess.

**Table 5:**
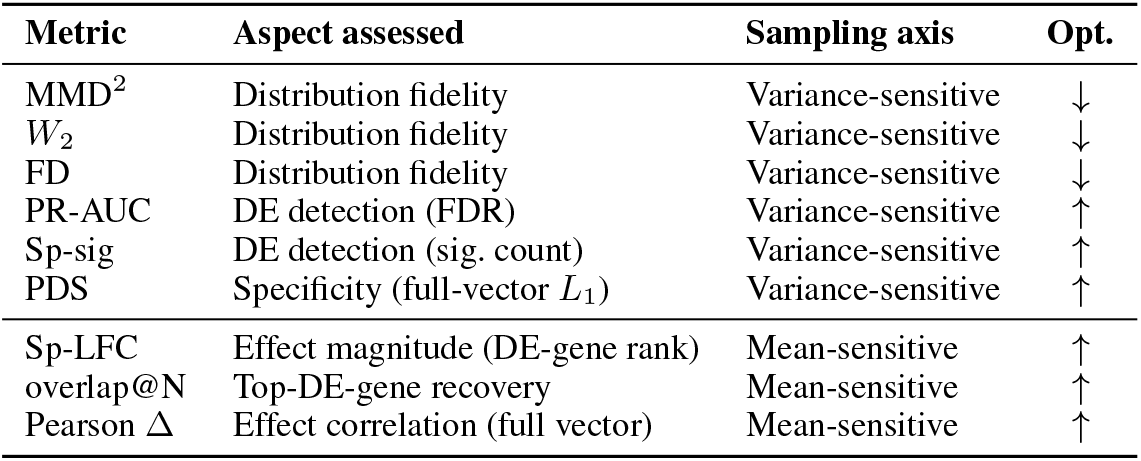
Summary of evaluation metrics. DE metrics share the same upstream pseudobulk and per-perturbation BH-FDR computation; distribution metrics operate in PCA-30 space fitted on ground-truth cells. The last column gives each metric’s inference-time sampling axis (Section 4.2): *variance-sensitive* metrics reward reproducing the within-perturbation distribution and full per-gene profile (favoring sampling heads), *mean-sensitive* metrics read a clean mean over strongly-DE genes (favoring deterministic heads). PDS is variance-sensitive despite being a pseudobulk metric: its *L*_1_ is dominated by the low-count bulk, unlike *Pearson* Δ, a correlation dominated by the strong-DE genes.

| Metric | Aspect assessed | Sampling axis | Opt. |
| --- | --- | --- | --- |
| MMD<sup>2</sup> | Distribution fidelity | Variance-sensitive | ↓ |
| $W_2$ | Distribution fidelity | Variance-sensitive | ↓ |
| FD | Distribution fidelity | Variance-sensitive | ↓ |
| PR-AUC | DE detection (FDR) | Variance-sensitive | ↑ |
| Sp-sig | DE detection (sig. count) | Variance-sensitive | ↑ |
| PDS | Specificity (full-vector $L_1$) | Variance-sensitive | ↑ |
| Sp-LFC | Effect magnitude (DE-gene rank) | Mean-sensitive | ↑ |
| overlap@N | Top-DE-gene recovery | Mean-sensitive | ↑ |
| Pearson $\Delta$ | Effect correlation (full vector) | Mean-sensitive | ↑ |

**Table 6:** Full prior × head ablation on Distribution metrics for Parse 1M and Replogle (HepG2). We add results from running 4-seed pooled mean ± SE runs. Bold = best in column within each dataset.

| Dataset | Prior | Head | $W_2$ ↓ | MMD<sup>2</sup> ↓ | FD ↓ |
| --- | --- | --- | --- | --- | --- |
| Parse 1M | AR | ce-quantile | 11.978±0.011 | **0.0039**±0.0001 | **1.78**±0.02 |
| | | nb-sample | 11.952±0.009 | 0.0081±0.0002 | 3.31±0.05 |
| | | hurdle | 12.045±0.007 | 0.0114±0.0003 | 5.00±0.10 |
| | | mse | 22.080±0.108 | 0.6462±0.0047 | 375.41±4.54 |
| | flow | ce-quantile | 12.243±0.008 | 0.0246±0.0002 | 11.06±0.15 |
| | | nb-sample | 12.600±0.007 | 0.0460±0.0005 | 20.48±0.24 |
| | | hurdle | 11.920±0.005 | 0.0073±0.0001 | 2.94±0.03 |
| | | mse | **11.893**±0.005 | 0.0111±0.0003 | 4.17±0.10 |
| | MDLM | ce-quantile | 11.997±0.004 | 0.0047±0.0001 | 2.13±0.04 |
| | | nb-sample | 11.969±0.007 | 0.0081±0.0001 | 3.50±0.06 |
| | | hurdle | 12.083±0.010 | 0.0129±0.0002 | 5.46±0.08 |
| | | mse | 22.152±0.073 | 0.6542±0.0031 | 378.10±2.81 |
| Replogle | AR | ce-quantile | **7.864**±0.010 | **0.0089**±0.0003 | 4.54±0.08 |
| | | nb-sample | 8.001±0.010 | 0.0095±0.0003 | 4.70±0.09 |
| | | hurdle | 7.900±0.010 | 0.0130±0.0004 | 5.47±0.11 |
| | | mse | 9.032±0.017 | 0.0987±0.0007 | 35.06±0.24 |
| | flow | ce-quantile | 8.049±0.008 | 0.0090±0.0001 | **4.34**±0.08 |
| | | nb-sample | 8.217±0.007 | 0.0192±0.0003 | 6.20±0.09 |
| | | hurdle | 7.885±0.005 | 0.0114±0.0002 | 4.93±0.06 |
| | | mse | 8.971±0.019 | 0.0783±0.0012 | 27.14±0.34 |
| | MDLM | ce-quantile | 8.051±0.020 | 0.0133±0.0004 | 5.36±0.15 |
| | | nb-sample | 8.121±0.014 | 0.0126±0.0004 | 5.29±0.10 |
| | | hurdle | 8.042±0.008 | 0.0172±0.0005 | 6.33±0.07 |
| | | mse | 9.091±0.021 | 0.1052±0.0011 | 35.70±0.38 |

**Table 7:**
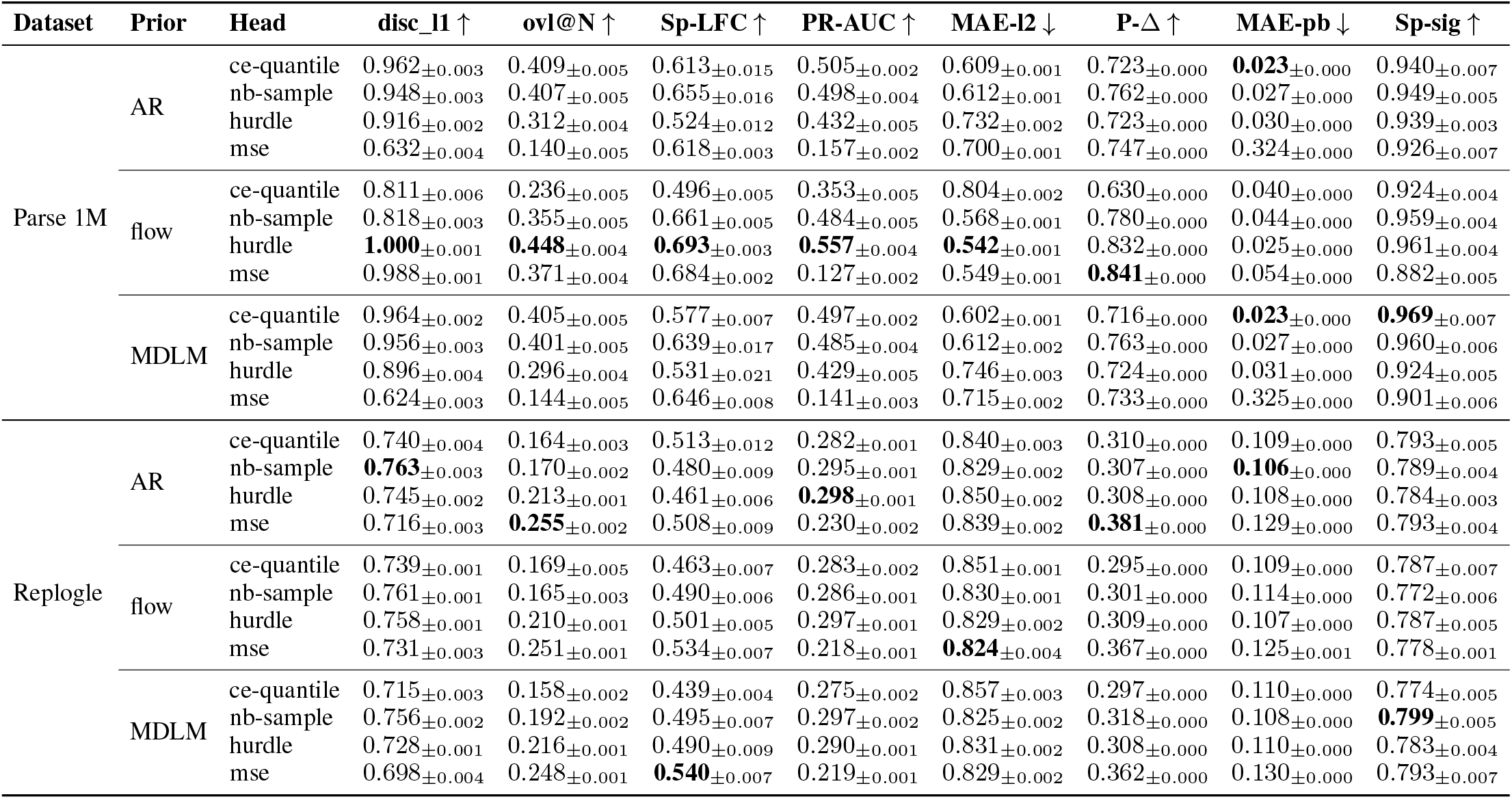
All prior × head ablations across cell-eval metrics.

## D Continous latent spaces

Upon the results presented in section C, we were curious if this followed continous latent spaces as well. The community has not released these versions among existing benchmarks, like ce-quantile, and hurdle with flow. Other ablations like mse and nb have been published across multiple studies. Here are the results for the full ablations on all the discrete and continous latent spaces. We also want to point out that while the flow model does better on the Parse 1M dataset - the same trend does not follow for the replogle dataset, and therefore it maybe just an artifact of the dataset. We hope to add further analysis in this study by expanding it to more datasets.

## Footnotes

1 https://sanjukta7.github.io/sclatents/

2 https://github.com/sanjukta7/sc-evae

